# Neural Mechanisms Underlying Approach and Avoidance Tendencies in Alcohol Use: An Electrophysiological Investigation

**DOI:** 10.1101/2025.06.14.658999

**Authors:** Adarsh K. Verma, Adith Deva Kumar, Usha Chivukula, Neeraj Kumar

## Abstract

**Background:** A growing body of research highlights the differential role of approach and avoidance tendencies toward alcohol cues in the development and maintenance of harmful drinking behavior. Some individuals involved in alcohol consumption show an automatic approach towards alcohol-related stimuli, whereas others demonstrate avoidance, suggesting the need to understand the neurocognitive mechanisms underlying these automatic tendencies. **Methods:** The current study employed an Alcohol Approach-Avoidance Task (A-AAT) with electroencephalography (EEG) to investigate neural responses among individuals with alcohol approach and avoidance tendencies. Alcohol group participants were categorized into approach and avoidance subgroups based on their behavioral tendencies following A-AAT administration. **Findings**: Results revealed significant attenuation in P3 and FN400 amplitudes at frontal and parietal sites, respectively, in the alcohol-approach participants compared to both the alcohol-avoidance and non-alcohol participants. These findings suggest weakened controlled cue processing and impaired stimulus-response conflict resolution in individuals with stronger approach tendencies. Notably, right prefrontal activity exhibited prominent differences between the approach and avoidance groups, highlighting its potential role in regulating automatic alcohol-related responses. **Implications**: The identified ERP markers provide clinical utility for assessing alcohol approach tendencies and monitoring the progress of intervention. Findings further emphasize the importance of individually tailored targeted interventions aimed at reducing harmful alcohol consumption behavior by altering alcohol approach tendencies.

## Introduction

Alcohol use is a complex behavior influenced by both automatic and controlled cognitive processes. While moderate alcohol consumption is socially acceptable, excessive and habitual use often leads to dependence and long-term negative consequences. Neurocognitive research highlights the role of automatic approach tendencies in driving alcohol consumption behavior (Ernst et al., 2014; McNaughton et al., 2016; C. E. Wiers et al., 2014). Individuals with strong alcohol approach tendencies exhibit a heightened, reflexive pull toward alcohol-related cues (Ernst et al., 2014; C. E. Wiers et al., 2014; R. W. Wiers et al., 2007). However, how avoidance tendencies might affect alcohol consumption is not clearly understood.

Approach and avoidance tendencies, rooted in Pavlovian reflexes, differ in their motivational underpinnings, attentional demands, and goal-directed behavioral responses (Moors, 2016; Najmi et al., 2010). A strong approach tendency has been linked to various maladaptive behaviors, such as addiction (C. E. Wiers et al., 2014; R. W. Wiers et al., 2002), whereas heightened avoidance tendencies have been associated with anxiety-related disorders, including obsessive-compulsive disorder (Krypotos, 2015; Najmi et al., 2010). Individuals with alcohol approach tendencies are more likely to develop and maintain alcohol dependence, struggle with relapse after abstinence, and exhibit heightened attentional biases toward alcohol-related cues (Eberl et al., 2013; Ernst et al., 2014; Van Gucht et al., 2008; C. E. Wiers et al., 2014). Paradoxically, individuals who exhibit alcohol avoidance tendencies may still engage in alcohol consumption despite their intention to abstain, particularly due to social influences, peer pressure, or negative emotional states (Kenney et al., 2018; McNaughton et al., 2016; Morris et al., 2020). Individuals with avoidance tendencies may experience an internal conflict between their will to withdraw from alcohol consumption and external pressures to drink, leading to cognitive dissonance that influences their drinking patterns (Jacobs et al., 2018; Kenney et al., 2018; Morris et al., 2020). Evidence from emotion and motivation research indicates that approach and avoidance tendencies differ across multiple levels, including unconscious stimulus evaluation, conscious interpretation, and behavioral responses (Corr, 2013; Hans Phaf et al., 2014; Warr et al., 2021). Unconscious evaluation of stimuli triggers emotional reactions, which form the basis of automatic predispositions (Bradley et al., 2001; Hans Phaf et al., 2014). Such affective responses play a critical role in guiding behavior, often resulting in an approach toward positive stimuli and avoidance of negative ones.

From a neurocognitive perspective, alcohol approach tendencies are associated with impairments in the neural correlates of cognitive control and goal-directed processing. Studies have shown that individuals with strong approach tendencies toward alcohol stimuli exhibit reduced activation in the dorsolateral prefrontal cortex (dLPFC) and medial prefrontal cortex (mPFC), regions critical for cognitive control, conflict monitoring, and decision-making (Ernst et al., 2014; Miller & Cohen, 2001). Electrophysiological evidence further suggests that high-risk drinkers show diminished N450 event-related potential (ERP) amplitudes in the prefrontal region, indicating deficits in stimulus-response conflict resolution (Cofresí, Kohen, et al., 2022; Imbir et al., 2018; Larson et al., 2014). Additionally, attenuation of the P3 ERP component at the parietal region has been observed in response to alcohol cues among heavy drinkers, reflecting deficits in controlled processing of stimulus and attentional regulation (Cofresí, Kohen, et al., 2022; Cofresí, Piasecki, et al., 2022; Cohen et al., 2002; Hajcak & Foti, 2020; Jurado-Barba et al., 2020). Reduced P3 amplitudes in individuals with alcohol dependence suggest impaired top-down cognitive control, making them more susceptible to automatic, cue-driven drinking behavior (Namkoong et al., 2004; Segalowitz et al., 2001).

The coexistence of both tendencies among individuals involved in alcohol consumption highlights the complexity of alcohol use and the necessity of distinguishing subgroups within this population in research and clinical practice. Despite growing evidence supporting the role of alcohol approach-avoidance tendencies in alcohol use, no study has explicitly examined how neural markers differ in individuals with alcohol avoidance tendencies, who may rely on different regulatory strategies to suppress alcohol-related impulses. Most studies have treated alcohol-consuming individuals as a homogeneous group, overlooking the distinction between individuals who exhibit impulsive approach tendencies toward alcohol cues and those who consume alcohol despite an avoidance tendency in specific contexts (Krypotos, 2015; Najmi et al., 2010). This homogeneous treatment of drinkers may conceal important neurocognitive differences that could inform more targeted intervention strategies.

The present study aims to investigate the differential neurocognitive profiles of individuals with opposing approach and avoidance tendencies using the Alcohol Approach-Avoidance Task (A-AAT) and electroencephalography (EEG). Specifically, we hypothesize that the individuals with alcohol avoidance tendencies and the non-alcohol group will show relatively enhanced P3 amplitudes (reflecting enhanced controlled processing of alcohol cues) compared to those with alcohol approach tendencies. Additionally, we expect that individuals with alcohol avoidance tendencies and non-alcoholic controls will exhibit enhanced FN400 ERP component (implying lower affective complexity and greater stimulus-response conflict resolution) compared to those with alcohol approach tendencies.

The findings indicate significantly attenuated centroparietal P3 and frontal FN400 amplitudes in the approach group when responding to alcohol cues, suggesting impaired cognitive processing and dominance of automatic behavior compared to the avoidance group. In contrast, the avoidance group shows no significant differences across alcohol and non-alcohol stimuli, revealing neural responses similar to the non-alcohol group.

## Method

### Participants

Thirty-nine right-handed male college students in the alcohol group (mean age = 22.8 years, SD = 3.45) and twenty in the non-alcohol group (mean age = 21.0 years, SD = 2.78) volunteered for the study. The participants provided written informed consent prior to their participation. All the participants had normal or corrected-to-normal vision and had no history of brain injury or neurological disorders. The alcohol group participants were screened using the Alcohol Use Disorders Identification Test (AUDIT; Saunders et al., 1993)LJ. Specifically, twenty-five individuals exhibited low-risk consumption patterns (scores ranging from 1 to 7), while twelve participants demonstrated harmful alcohol consumption (scores between 8 and 14). Two participants scored above 14, indicating a likelihood of developing alcohol dependence. At the time of the study, none of the participants were diagnosed with alcohol use disorder (AUD). All the participants were instructed in advance to abstain from alcohol use for at least 24 hours prior to the experiment. A verbal confirmation of compliance with the abstinence requirement was obtained from each participant before the session began. The study was conducted in accordance with the Declaration of Helsinki, with approval from the Institutional Ethics Committee.

### Trial Procedure

The Alcohol Approach Avoidance Task (A-AAT; Korucuoglu et al., 2016; Rinck & Becker, 2007; C. E. Wiers et al., 2014)LJ was adapted and administered using Psychtoolbox in MATLAB (The MathWorks, Natick, MA). Forty picture stimuli were used: twenty alcoholic beverage images and twenty non-alcoholic images. The non-alcoholic stimuli included images of water and juice, serving as neutral controls. The alcoholic and non-alcoholic pictures were matched for valence and arousal using the Self-Assessment Manikin framework (SAM; Bradley & Lang, 1994)LJwith an independent sample of eight low-risk drinkers.

The study was structured into two experimental blocks. In the initial block, a total of one hundred sixty trials were conducted, comprising eighty randomly presented trials for both alcohol-related images and neutral images. The participants’ responses were recorded via joystick movement, where they would pull the joystick towards their body in response to alcohol images and push away from their body in response to neutral images, simulating the alcohol approach/avoidance condition. In the second block, the movements were reversed, i.e., alcohol stimuli were pushed, and non-alcohol stimuli were pulled. A washout block was administered between the two experimental blocks, during which participants were instructed to pull images of pens and push images of screwdrivers. This procedure aimed to eliminate use-dependent behavioral characteristics that might carry over from the previous block. Pull and push movements were programmed to produce zoom-in and zoom-out effects, respectively, with the magnitude of the effect proportional to the joystick’s displacement along the y-axis. At the stimulus onset, the image appears at the center, occupying 50% of the display area. A maximum pull (+1 on the y-axis) enlarges the image to fill the entire screen, while a maximum push (−1 on the y-axis) reduces the image size until it completely disappears.

Each trial started with a 500 ms fixation period, followed by the onset of the stimulus image. The response window allowed for a maximum reaction time of 3000 ms, within which participants could respond to the stimulus. Feedback was provided as a red cross for incorrect trials, with a duration of 500 ms. Before each trial, a minimum time constraint was established to prevent unintentional movements and false reactions, requiring the participant to maintain a neutral joystick position for at least 200 ms before responding. Any violation of this (i.e., early movement) would reset the 200-ms timer, delaying the onset of the stimulus.

### EEG Recording and Preprocessing

The EEG data were recorded using the Brain Products actiCHamp Plus system (Brain Products GmbH, Munich, Germany), consisting of 32 active Ag/AgCl electrodes positioned according to the standard 10-20 montage. The sampling rate was set at 5000 Hz, and electrode impedance was maintained below 10 kΩ. Data acquisition was referenced to the Cz electrode.

The recorded EEG signals were then pre-processed using the EEGLAB toolbox (Delorme & Makeig, 2004) and custom-written MATLAB scripts tailored to specific requirements. During pre-processing, the data were downsampled to 1000 Hz. The initial bandpass filter with a frequency range of 0.1-30 Hz was applied, followed by averaged re-referencing. The EEG data were then subjected to event-related epoching from −500 ms to 1000 ms relative to stimulus (alcohol/non-alcohol image) onset, with baseline correction using a window of −200 to 0 ms, followed by epochs binning for the statistical analysis. Manual artifact rejection was employed to identify and remove artifactual trials or anomalies. Eyeblinks and muscle artifacts were removed from the EEG signals with the help of independent component analysis (ICA using EEGLAB’s runica algorithm).

A bandpass filter of 0.2 to 10 Hz was employed on the preprocessed data to extract P3 and FN400 components based on established methodologies, (Zhang et al., 2024b, 2024a). The FN400 mean amplitude was calculated within a 400–600 ms time window at frontal electrodes F3, F4, and Fz. The P3 mean amplitude was computed within a 300–600 ms time window at the Pz electrode. The choice of time windows was also guided by visual inspection, allowing for the identification of optimal parameters for each component.

### Statistical Analysis

In the AAT task, each trial’s response time (RT) was computed as the difference between the stimulus onset and the moment the joystick reached its maximum position along the y-axis. For each image type (alcoholic and non-alcoholic), an automatic tendency score was calculated by subtracting median push reaction times from pull reaction times. Then, the Alcohol Approach Index (AAI) was computed for each participant by subtracting the score for non-alcoholic images from the score for alcoholic images. This provided a standardized measure of the approach/avoidance tendency towards alcoholic stimuli. A positive AAI score indicated that individuals exhibited an approach toward alcohol-related images, suggesting a greater inclination to approach or seek out alcohol-related stimuli. Conversely, a negative AAI score suggested that individuals show an avoidance tendency towards alcohol-related images, implying a reduced interest in approaching or seeking out alcohol-related stimuli. Based on these scores, participants in both the alcohol and non-alcohol groups were initially categorized into approach and avoidance subgroups to examine potential differential neural responses.

For ERP analysis, stimulus-locked mean ERP amplitudes were calculated at specific electrode sites by averaging binned epochs for each condition (combinations of movement direction (pull, push) and stimulus category (alcohol, non-alcohol)). Wilcoxon rank-sum tests were first conducted to assess whether ERP responses significantly differed between the alcohol and non-alcohol groups. Additionally, ERP responses were compared between individuals categorized as having approach or avoidance tendencies within both the alcohol and non-alcohol groups to determine whether these behavioral dispositions corresponded with distinct neural patterns. As no statistically significant differences were observed between the approach and avoidance subgroups within the non-alcohol group, these participants were pooled into a single group for subsequent analyses.

A non-parametric aligned ranks transformation ANOVA (Wobbrock et al., 2011) was conducted to examine the effects of group (alcohol approach, alcohol avoidance, non-alcohol), stimulus type (alcohol, non-alcohol), and joystick movement (pull, push) on ERP amplitudes, as the data were not normally distributed and/or the assumption of homogeneity of variance was violated.

Wilcoxon pairwise comparisons were performed for significant ANOVA results, with p-values adjusted using the Benjamini-Hochberg (BH) correction. Effect sizes (r) were computed for pairwise comparisons. Additionally, Spearman correlation analyses were conducted between AAI scores and the difference in ERP mean amplitudes (alcohol minus non-alcohol stimulus) for each electrode site. The use of ERP subtraction scores aligns with the computation of the AAI score, which reflects the difference in automatic tendencies toward alcohol versus non-alcohol stimuli. Correlating ERP difference scores with the behaviorally derived difference score allows conceptual consistency in assessing how neural responses relate to individual variations.

All statistical analyses were conducted in R (version 4.3.1). The ARTool package was used for non-parametric ANOVA. (Wobbrock et al., 2011), and the wilcox_effsize function was used to compute effect sizes (Tomczak & Tomczak, 2014).

## Results

The current study aimed at exploring neurocognitive differences in stimulus evaluation and conflict resolution among alcohol subgroups with automatic approach and avoidance tendencies toward alcohol, as well as a non-alcohol group, using ERP measures. Alcohol and non-alcohol users were sub-classified into the approach and avoidance groups based on their Approach-Avoidance Index (AAI). Within the alcohol group, 20 participants exhibited an approach tendency, as indicated by a positive AAI score (0.085 ± 0.046), meaning they responded faster when pulling alcohol-related images than when pushing them, suggesting a predisposition to approach alcohol cues. Conversely, 19 individuals displayed an avoidance tendency, characterized by a negative AAI score (−0.088 ± 0.060), indicating they were faster at pushing than pulling alcohol-related images, reflecting an avoidance response. In the non-alcohol group, 10 individuals showed a behavioral approach tendency toward alcohol (0.053 ± 0.024), and 10 showed avoidance (−0.081 ± 0.029).

### ERP Differences Across Groups and Automatic Behavioral Tendencies

Preliminary analyses revealed significant differences in event-related potential (ERP) responses between the alcohol and non-alcohol groups. Specifically, the alcohol group exhibited reduced P3 mean amplitudes at the Pz electrode (median difference = −2.668; W = 4117, p_adj_ < 0.001, r = 0.278), and reduced FN400 ERP at F4 (median difference = 1.352; W = 7898, p_adj_ = 0.001, r = 0.217) and Fz sites (median difference = 2.279; W = 8495, p_adj_ < 0.001, r = 0.296). No significant difference was observed for FN400 amplitude at F3 (median difference = 1.621; W = 7314, p_adj_ = 0.092).

Within the non-alcohol group, participants classified with approach versus avoidance tendencies (based on AAI scores) did not show significant differences in ERP amplitudes across electrodes, i.e., P3 mean amplitude at Pz (median difference = 0.741; W = 921, p_adj_ = 0.248) and for FN400 mean amplitude at F4 (median difference = 0.778; W = 850, p_adj_ = 0.636) or Fz (median difference = −1.715; W = 747, p_adj_ = 0.615). Given the lack of ERP differences between behavioral subgroups in the non-alcohol group, these individuals were treated as a single reference group in the subsequent analyses.

In contrast, significant ERP differences emerged within the alcohol group based on automatic behavioral tendencies. Participants with an approach bias exhibited lower ERP amplitudes (P3 ERP at Pz, median difference = −0.345, W = 2240, p_adj_ = 0.007, r = 0.227; and FN400 at F4, median difference = 0.761, W = 4118, p_adj_ < 0.001, r = 0.306; and Fz, median difference = 0.982, W = 4133, p_adj_ < 0.001, r = 0.310), relative to avoidance-biased individuals. These initial results justify subsequent analyses focusing on differences in ERP responses across three groups: non-alcohol participants (approach and avoidance group combined), alcohol-approach individuals, and alcohol-avoidance individuals with respect to stimulus type and movement direction.

### Influence of Automatic Tendencies and Stimulus Type on Controlled Processing of Stimulus (P3 ERP)

Participants showed differential P3 ERP activity at the Pz electrode dependent on the automatic tendencies and stimuli (Figure 2A), with a significant group × stimulus type interaction (F_(2,168)_ = 30.659, p < 0.01). Further, pairwise comparisons showed that while responding to alcohol cues, the approach group exhibited significantly lower P3 amplitudes (Figure 3A) compared to the avoidance group (median difference = −2.161; W = 100, p_adj_ = 0.036, r = 0.387) and the non-alcohol group (median difference = −4.434; W = 89, p_adj_ = 0.013, r = 0.509). However, no significant difference was observed between the avoidance and non-alcohol groups (median difference = −2.273; W = 153, p_adj_ = 0.458). Additionally, within-group comparisons revealed a significant difference in P3 amplitudes between alcohol and non-alcohol stimuli only in the approach group (median difference = −1.975; W = 6, p_adj_ = 0.002, r = 0.801), whereas no such effect was found in the alcohol avoidance (median difference = 0.350; W = 111, p_adj_ = 0.605) or non-alcohol groups (median difference = −0.169; W = 88, p_adj_ = 0.605).

**Figure 1.**
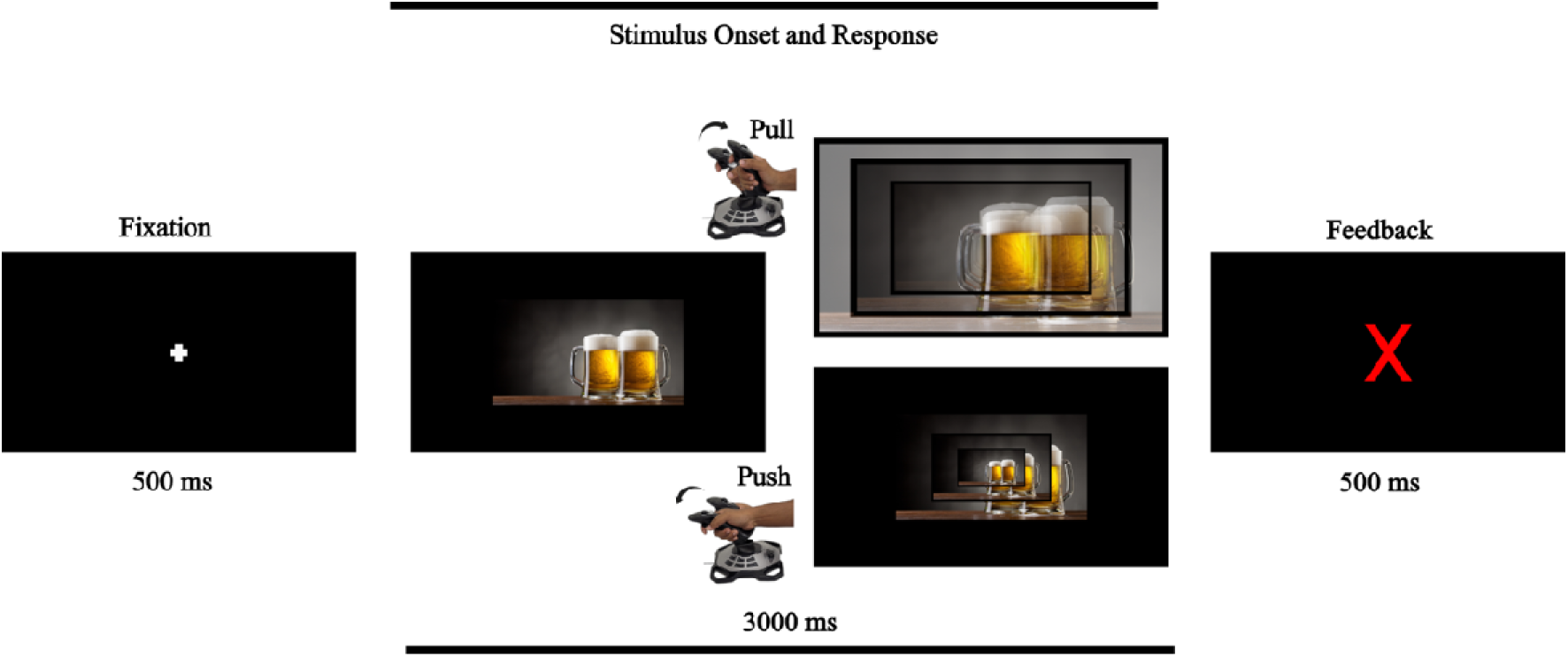
Schematic representation of an AAT trial. In the task, the images/stimuli are presented at the center of the screen, where participants pull/push each image using the joystick depending on the instruction (more details in the task/stimuli section). The trial starts with a fixation (+) for 500 ms, followed by the stimulus presentation within a response time window of 3000 ms, where the response results in zooming in/out of the stimulus depending on pull/push movement, respectively. This screen is followed by the cross (X) symbol in case of wrong movement for 500 ms.

**Figure 2.**
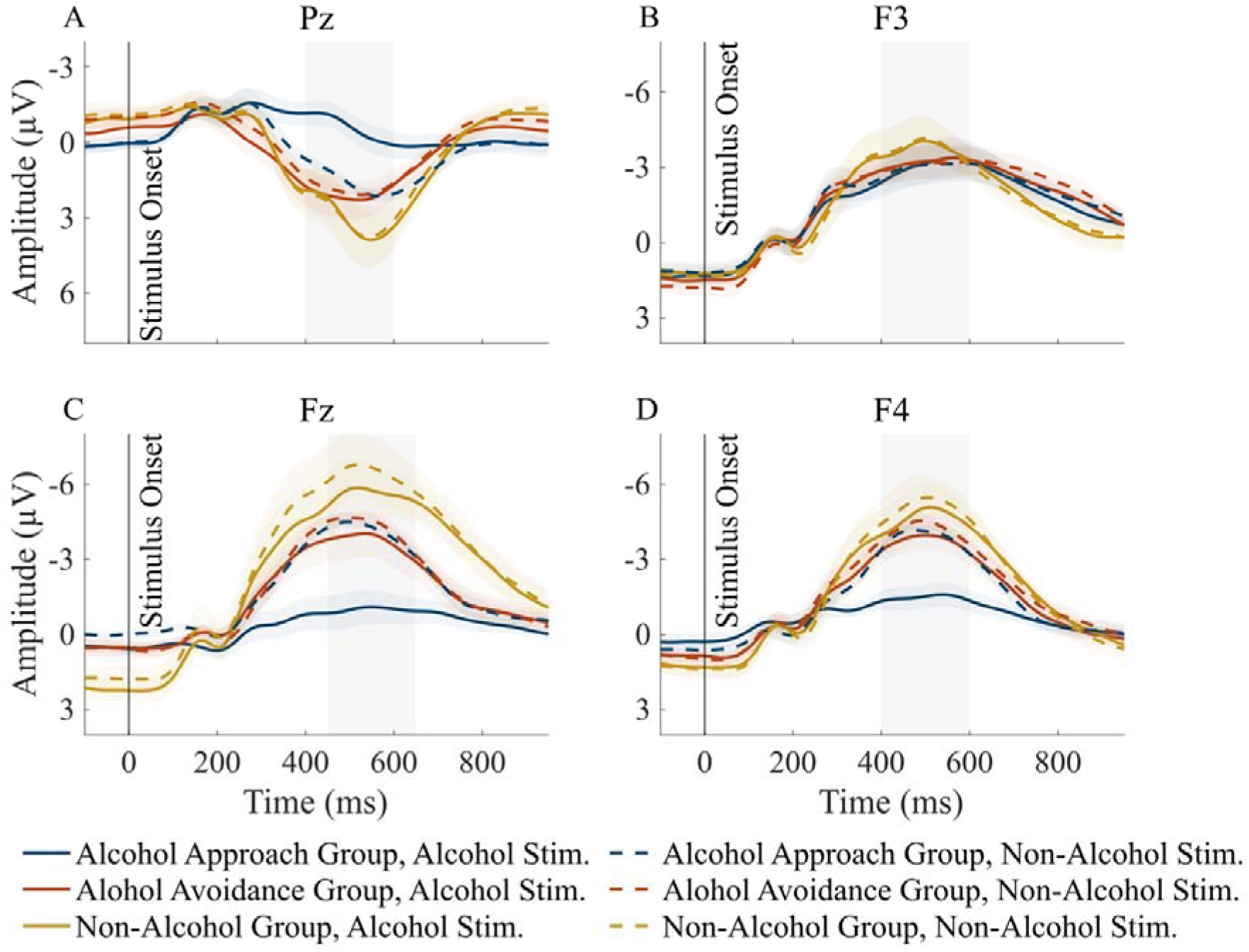
The participants with approach tendencies exhibit reduced neural responses to alcohol stimuli compared to those with avoidance tendencies. Panel A–D show the averaged ERP amplitude for the Approach (blue color), Avoidance (red color), and Non-Alcohol (yellow color) groups in response to alcohol (solid lines) and non-alcohol (dashed lines) stimuli at the Pz (Panel A), F3 (Panel B), Fz (Panel C), and F4 (Panel D) electrodes. The highlighted areas represent the measurement windows that quantify the P3 ERP at Pz and FN400 at Fz, F3, and F4.

**Figure 3.**
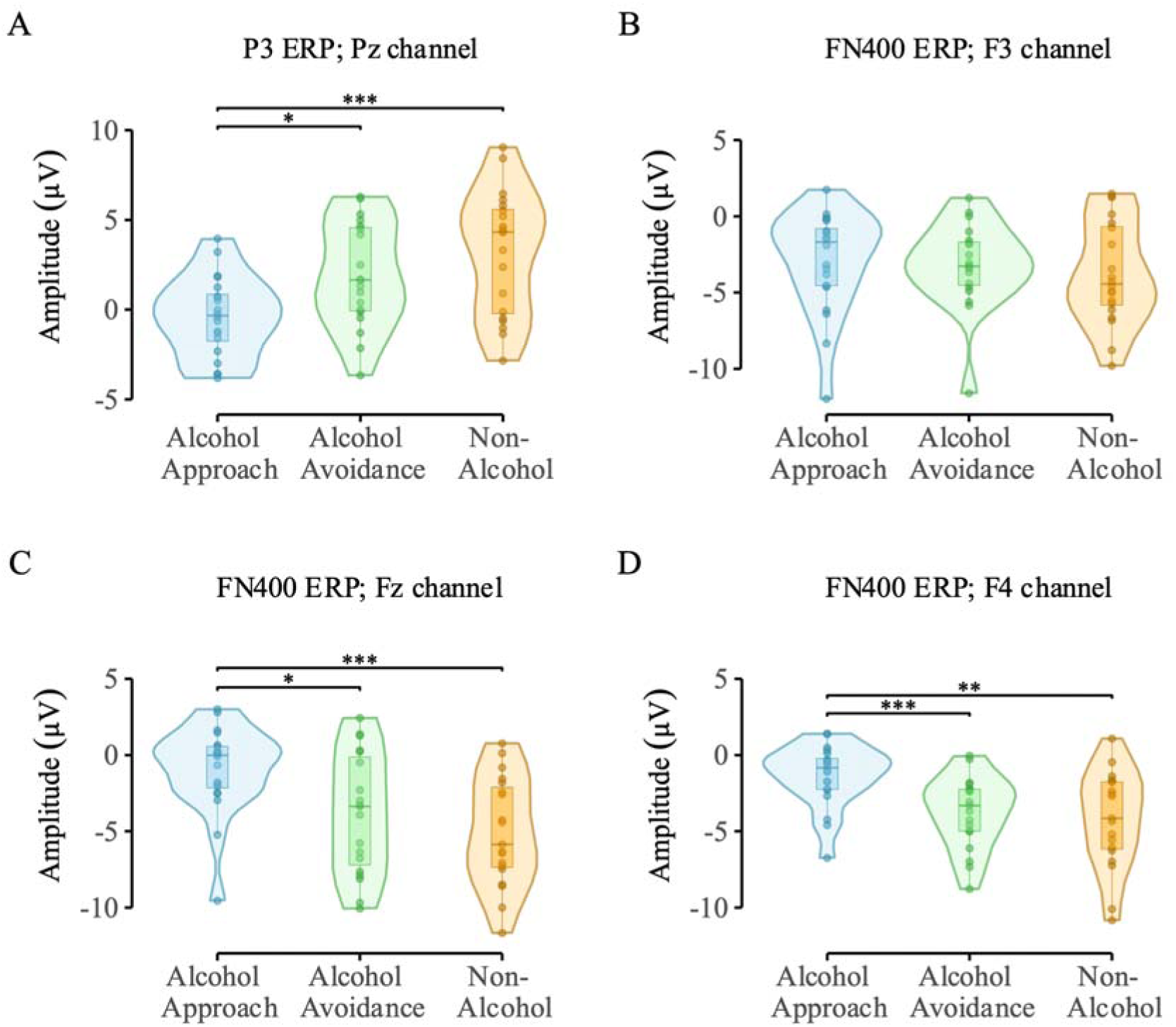
Participants in the approach group exhibited significantly attenuated ERPs compared to those in the avoidance and non-alcohol groups. Panels A–D display P3/FN400 ERP distributions in response to alcohol stimuli at the Pz (Panel A), F3 (Panel B), Fz (Panel C), and F4 (Panel D) electrodes. * p < 0.05; ** p < 0.01; *** p < 0.001

The observed group and stimuli effects on P3 ERP amplitude were not found to be influenced by the movement direction, as indicated by a non-significant group × stimulus type × movement interaction (F_(2,168)_ = 0.939, p = 0.393), suggesting joystick movement (pull, push) had minimal impact on controlled stimulus processing.

These results suggest that an automatic approach tendency toward alcohol is associated with diminished controlled processing of alcohol-related stimuli, as reflected in reduced P3 amplitudes. In contrast, avoidance tendencies do not appear to influence the engagement of controlled evaluation processes, as evidenced by the similar P3 amplitude patterns in the approach and avoidance groups.

### Effect of Automating Tendencies and Stimulus Type on Conflict Processing and Resolution (FN400 ERP)

At the frontal region, participants with approach tendency showed varied neural responses (FN400 amplitudes) compared to the participants with avoidance tendency for different stimulus types (Figure 2, 3; Panels B, C, and D). ANOVA shows significant group × stimulus type interaction for FN400 amplitude at F4 (F_(2,168)_ = 38.670, p < 0.01) and Fz (F_(2,168)_ = 21.290, p < 0.01), but not at F3 (F_(2,168)_ = 1.284, p = 0.280), suggesting a lateralized rather than a uniform effect of automatic tendencies and stimulus type at the frontal sites. Pairwise comparisons highlight significant differences in FN400 amplitude between the approach and avoidance groups for alcohol stimuli (Figure 2, Panels C and D) at F4 (median difference = 2.551; W = 302, p_adj_ = 0.005, r = 0.513) and Fz (median difference = 3.439; W = 281, p_adj_ = 0.025, r = 0.378). Similarly, the approach group showed significantly different FN400 amplitudes from the non-alcohol group at F4 (median difference = 3.886; W = 321, p_adj_ = 0.005, r = 0.522) and Fz (median difference = 5.325; W = 340, p_adj_ = 0.001, r = 0.556).

However, no significant differences were found between the avoidance and non-alcohol groups at F4 (median difference = 1.334; W = 222, p_adj_ = 0.376) or Fz (median difference = 1.886; W = 240, p_adj_ = 0.211), suggesting that these two groups may engage similar neural mechanisms in effectively resolving affective conflicts when encountering alcohol stimuli in contrast to the approach group, which exhibited significantly attenuated FN400 ERP pattern. This inference is further supported by within-group comparisons, where the approach group exhibited significantly reduced FN400 amplitudes when responding to alcohol versus non-alcohol stimuli at both F4 (median difference = 2.689; W = 209, p_adj_ = 0.001, r = 0.860) and Fz (median difference = 3.481; W = 192, p_adj_ = 0.005, r = 0.735). In contrast, the avoidance group did not have a significant difference in FN400 amplitude across stimulus types at either F4 (median difference = 0.650; W = 124, p_adj_ = 0.375) or Fz (median difference = 0.867; W = 112, p_adj_ = 0.507). However, the non-alcohol group exhibited a significant difference in FN400 amplitude at Fz (median difference = 0.662; W = 189, p_adj_ = 0.005, r = 0.743) but not at F4 (median difference = 0.463; W = 153, p_adj_ = 0.171).

Similar to P3 ERP, movement direction during the A-AAT task did not significantly interact with group and stimulus type on FN400 mean amplitude at F4 (F(2,168) = 0.540, p = 0.584) and Fz (F(2,168) = 0.448, p = 0.640). This suggests that approach tendencies and stimulus type primarily influenced FN400 amplitude modulation rather than the action.

Overall, these results indicate that individuals with an alcohol approach tendency exhibit attenuated FN400 responses to alcohol stimuli at both F4 and Fz, reflecting compromised response conflict resolution when alcohol-related cues elicit competing prepotent responses. In contrast, the alcohol avoidance and non-alcohol groups do not demonstrate this differentiation, implying that affective complexity in alcohol-related processing is unique to individuals with an automatic approach tendency.

### Heightened Alcohol Approach Tendencies Associated with Attenuated P3 and FN400 ERPs in Response to Alcohol Stimuli

A correlation analysis assessed whether observed ERP response patterns were linked to the corresponding behavioral response. Specifically, ERP difference scores (mean amplitude for alcohol stimuli minus non-alcohol stimuli) were correlated with AAI scores for conceptual consistency, as both reflect alcohol-related biases. Between AAI and P3 amplitude difference at the Pz electrode, a significant negative correlation (ρ = - 0.40, p = 0.003) was observed. It suggests that stronger alcohol approach tendencies were associated with diminished P3 responses to alcohol stimuli, resulting in a more negative difference score (alcohol – non-alcohol). As AAI scores become negative, reflecting avoidance tendencies, this difference diminishes, implying similar engagement with alcohol and non-alcohol stimuli during controlled processing.

The FN400 ERP difference at the F4 electrode was significantly correlated with the AAI, showing a positive association (ρ = 0.41, p = 0.003). This indicates that individuals with stronger approach tendencies toward alcohol (higher AAI scores) exhibited attenuated (less negative) FN400 responses to alcohol stimuli relative to non-alcohol stimuli, resulting in greater positive difference scores. Conversely, as AAI scores decrease toward zero or negative values (reflecting avoidance tendencies), the FN400 difference also reduces, indicating more similar neural responses to both alcohol and non-alcohol stimuli (Figures 3 and 4).

Additionally, strong correlations were found between different electrode sites. F4 and Fz were positively correlated (ρ=0.52, p<0.001), while both F4-Pz (ρ=−0.56, p<0.001) and Fz-Pz (ρ=−0.65, p<0.001) showed negative correlations. These findings reflect the inverse relationship between frontal (F4, Fz) and parietal (Pz) responses, where higher P3 amplitude (more positive) at the Pz electrode and higher FN400 amplitude (more negative) at the F4 and Fz electrodes indicate better processing. Attenuated amplitudes (closer to zero) in the alcohol approach group while responding to alcohol stimuli suggest weaker neural responses, which may indicate automatic, less efficient processing of alcohol cues.

## Discussion

Contemporary models of addiction, particularly dual-process frameworks, emphasize the dynamic interplay between automatic motivational processes (e.g., approach biases) and deliberate cognitive control as key mechanisms underlying substance use behaviors (Lindgren et al., 2019; R. W. Wiers et al., 2007). However, these models often treat alcohol users as a homogeneous group, potentially obscuring important individual differences in automatic tendencies, which may lead to divergent patterns of neural engagement (e.g., cognitive control) and behavioral outcomes (e.g., impulsive drinking, cue-induced relapse, or successful self-regulation).

The present study categorized individuals based on their approach-avoidance profile, aiming to identify distinct neurocognitive patterns that may be masked when alcohol users are considered as a single group. Group differences revealed that individuals with an approach bias exhibit deficits in controlled cognitive processing and affective conflict resolution, as reflected in attenuated P3 and FN400 amplitudes, relative to alcohol avoidance and non-alcohol groups. Specifically, the reduced FN400 negativity in the alcohol approach group indicates impaired conflict resolution, likely driven by the dominance of automatic, impulsive reactions toward prepotent responses in the presence of alcohol stimuli. Such responses may evoke emotional or motivational conflict, consistent with previous findings (Imbir et al., 2018; Lepock et al., 2024). Attenuation in P3 amplitude among approach-biased individuals suggests reduced cognitive control and diminished top-down attentional regulation during alcohol cue processing. These neural deficits were not evident in the alcohol-avoidance or non-alcohol groups, which demonstrated relatively preserved conflict resolution and regulatory engagement.

The negative correlation between approach-avoidance index (AAI) and P3 amplitude difference at Pz site further demonstrates that increased approach biases are associated with weaker controlled processing in the parietal cortex, possibly reinforcing habitual alcohol-seeking behaviors, contributing to impulsive responses (Cohen et al., 2002; Elmasian et al., 1982; Emmerson et al., 1987; Euser et al., 2012; Jones et al., 2013; Jurado-Barba et al., 2020; Plawecki et al., 2018). The correlation between the AAI and FN400 amplitude difference at the right frontal location aids group-level findings, indicating that alcohol approach biases are linked to weaker conflict processing. Reduced affective conflict resolution during alcohol cue processing leads to heightened automatic approach behaviors, making it more difficult to disengage from substance-related stimuli (Boog & Tibboel, 2023; Christiansen et al., 2012; Moors, 2016; Pan et al., 2020). This pattern aligns with inhibitory control models, which emphasize the role of frontal regions in suppressing prepotent responses to salient cues (Brown et al., 2015; Hasher & Campbell, 2020; Ridderinkhof et al., 2004).

The pattern of ERP amplitude variations across parietal (Pz) and frontal (F4, Fz) sites highlights the interaction between cognitive control and affective conflict resolution during alcohol cue processing (Cofresí, Kohen, et al., 2022; Oscar-Berman & Marinković, 2007; Tapert et al., 2001). Negative correlations between P3 and FN400 components suggest that reduced attentional engagement is associated with impaired regulation of automatic responses. Positive correlations between frontal sites (F4 and Fz) reflect the coordinated recruitment of neural resources to detect and resolve response conflicts (Imbir et al., 2018; Larson et al., 2014; Namkoong et al., 2004). Collectively, these findings indicate that weaker ERP responses across parietal and frontal regions may underlie difficulties in suppressing automatic approach tendencies toward alcohol cues.

These findings extend previous research by demonstrating that categorizing drinkers based on their implicit motivational tendencies can uncover neurocognitive profiles, such as attenuated neural responses in the alcohol approach group compared to the avoidance group, that may not be evident when treating drinkers as a uniform group. Failing to distinguish these subgroups may lead to an oversimplified understanding of the involved mechanisms underlying alcohol-related behavior.

The results also contribute to dual-process models of addiction by clarifying the neurophysiological underpinnings of automatic and controlled processing in alcohol use (Lindgren et al., 2019). Particularly, the findings suggest that maladaptive alcohol consumption behaviors emerge from an imbalance between automatic approach tendencies (not avoidance tendencies) and controlled regulatory mechanisms. Additionally, this study addresses a key limitation in existing theories, i.e., overlooking individual differences in motivational directionality (Lannoy et al., 2014; Lindgren et al., 2019). By differentiating between individuals with alcohol-approach versus alcohol-avoidance tendencies, the study highlights that not all at-risk individuals exhibit the same automatic inclinations.

This categorical approach also holds practical implications. Categorizing drinkers based on implicit tendencies could provide more accurate behavior predictions, improve intervention targeting, and ultimately design tailored treatment strategies aligned with individual predispositional profiles. Furthermore, it may help identify relapse vulnerability pathways: individuals with strong approach biases may be more prone to cue-induced relapse due to diminished regulatory control, whereas avoidance-oriented individuals, though more behaviorally regulated, may experience heightened internal conflict, necessitating alternative forms of support (Eberl et al., 2013; Schlauch et al., 2012). The present study provides a foundation for more nuanced, effective treatment approaches by capturing this functional heterogeneity.

Nevertheless, certain limitations should be addressed in future research, particularly the generalizability of these findings to clinical populations, as the current sample does not fully represent individuals with alcohol dependence. From an interventional perspective, further studies should evaluate behavioral and neural interventions targeting alcohol approach tendencies. Additionally, investigating the functional connectivity underlying strong automatic alcohol approach tendencies may provide deeper insights into the mechanisms driving these effects.

In conclusion, the present study offers important insights into the distinct neurocognitive underpinnings underlying automatic approach and avoidance tendencies toward alcohol cues. This study highlights functionally meaningful differences in neural processing by moving beyond traditional unidimensional analyses and categorizing individuals based on their implicit motivational orientation. Individuals with an alcohol avoidance tendency exhibited enhanced neural activity, closely resembling the neural responses of the non-alcohol group. In contrast, the alcohol approach group shows deficits in these regulatory processes. These findings underscore the critical need to account for motivational directionality when studying alcohol-related behavior. By categorizing individuals into approach- and avoidance-oriented subgroups, researchers and clinicians can more accurately predict behavioral outcomes, tailor cognitive interventions, and develop more targeted strategies to mitigate relapse risk.

## CONFLICT OF INTEREST

None

## ACKNOWLEDGEMENT

We thank the Sensorimotor Plasticity Lab at the Indian Institute of Technology (IIT) Hyderabad for providing the necessary resources and support for this work.

## DATA AVAILABILITY

The datasets used and/or analyzed during the current study are available from the corresponding author upon reasonable request.

## FUNDING

This work was supported by the Ministry of Education, Government of India, through the Prime Minister’s Research Fellow (PMRF) scheme granted to AKV.

## Notes

### Competing Interest Statement

The authors have declared no competing interest.

